# Phylogenomic description of three novel species of the *Microbulbifer* genus, phylum Pseudomonadota, isolated from marine sponges and corals

**DOI:** 10.64898/2026.06.10.731415

**Authors:** Yifan Tang, Ashley P. Track, Natalie A. Miller, Paige Mandelare-Ruiz, Valerie J. Paul, Konstantinos T. Konstantinidis, Vinayak Agarwal

## Abstract

Understudied bacterial genera present a dynamic phylogenetic landscape and opportunities for discovering new taxa as more strains are isolated and genomic data is added. Here, through phylogenomic analysis, we describe three novel species of the globally distributed cosmopolitan marine bacterial genus *Microbulbifer*. This genus is ubiquitous in saltwater microbiomes and is a validated source of biodegradation enzymes as well as high value small molecule natural products. Average nucleotide identity (ANI) to the closest known species, *Microbulbifer variabilis* ATCC 700307^T^, was less than 88.4% for all three novel species. Isolates of the three novel species, designated as PAAF003^T^ (T = type strain), ZKSA006^T^, and SSSA003^T^ were imaged to reveal their phormological characteristics. Based on phylogenetic data, strains PAAF003^T^, ZKSA006^T^, and SSSA003^T^ represent three new species of the genus *Microbulbifer*, for which the names *Microbulbifer maximicatervae* sp. nov., *Microbulbifer regidiadema* sp. nov., and *Microbulbifer mixtoriginis* sp. nov. are proposed, respectively, under the SeqCode. We also reconstructed a robust phylogeny of available *Microbulbifer* genomes, which should faciliatate future isolation and strain description studies.

---

The genus *Microbulbifer* consititutes halophilic, rod-shaped, aerobic Gram-negative bacteria frequently isolated from marine holobionts such as sponges, algae, and corals, as well as from marine and mangrove sediments. These bacteria are known for their ability to degrade complex polysaccharides [1]. In addition, *Microbulbifer* strains produce structurally elaborate secondary metabolites—colloquially referred to as natural products—with established pharmaceutical utility [2-6]. Together with their clinically useful bioactivities, the structural complexity of *Microbulbifer*-derived natural products has inspired chemical synthesis innovation as well as discovery of novel natural product biosynthetic enzymology [7-9]. The inventory for the *Microbulbifer* genus has expanded to 39 species, according to the List of Prokaryotic names with Standing in Nomenclature (LPSN) with over 118 genomes now available. Given the ubiquity of this genus in marine and mangrove environments and the rapidly increasing strain and genome inventory, it is timely to conduct a comprehensive phylogenomic re-examination of the *Microbulbifer* genus. Furthermore, the genomic potential for natural product biosynthesis in *Microbulbifer* is species-specific, as we have described previously [10]. Hence, taxonomic benchmarking will also facilitate strain prioritization for future chemical discovery efforts from the *Microbulbifer* genus.

We have previously reported a large-scale bacterial cultivation effort from marine sponges and corals collected in the Florida Keys [11]. Among the 42 *Microbulbifer* strains isolated, we detected the presence of novel species based on currently used genome-based criteria [10]. The analysis used to justify the categorization of these strains into new genomic species—also called genomospecies—was based on comparison to known *Microbulbifer* type-strains. Here, we conducted a more comprehensive analysis using *all* publicly available 16S rRNA and whole genome sequences to robustly place the previously identified genomospecies within the context of known *Microbulbifer* phylogeny. We also conducted a thorough meta-analysis of geography and isolation source of the updated *Microbulbifer* strain inventory. In addition to naming three novel *Microbulbifer* species, our results indicate a marginal correlation between species diversity and the isolation source.

From the 42 *Microbulbifer* strains that we had isolated from marine sponges and corals, the presence of three novel genomic species was detected, which we had designated as GS-I–III (GS:genomospecies) [10]. The genomes for all 42 strains were sequenced, assembled, and are now available in the public databases. These genomes were queried alongside 76 additional whole genome assemblies of *Microbulbifer* strains available in GenBank and 7 additional strains for which only the 16S rRNA gene sequences were available (Table S1). Together with three more deep-branching sequences, the 16S rRNA gene sequences were retrieved to construct a maximum-likelihood phylogenetic tree (Figure S1). The resulting tree demonstrated distinct clades for GS-I–III (Figure S2), consistent with our earlier study [10]. These observations allowed us to posit that GS-I–III could represent new *Microbulbifer* species.

To further assess the genomic distinctness of GS-I–III, we determined the whole genome average nucleotide identity (ANI) values among the 118 *Microbulbifer* strains for which genome assemblies are available (42 sequenced as part of our previous study [10], and 76 available in GenBank) (Figure 1, Figure S3–S5, Supplementary Dataset D1). We previously demonstrated that as compared to 16S rRNA sequence-based phylogeny, ANI provides a higher resolution for species designation within the *Microbulbifer* genus [12]. Congruent with the phylogenetic tree for the 16S rRNA sequences, the ANI analysis also showed that genomes of GS-I–III are organizated into distinct clades (Figure 1). For GS-I– III, the closest type strain to all three genomic species was *Microbulbifer variabilis* ATCC 700307^T^ with an ANI values across strains ranging between 84.1% and 88.4% [13]. Based on all available *Microbulbifer* phylogenetic data to date, we posit that GS-I–III are novel species of the genus *Microbulbifer* and propose the following strain designations under the SeqCode nomenclature code [14].

**Figure 1.**
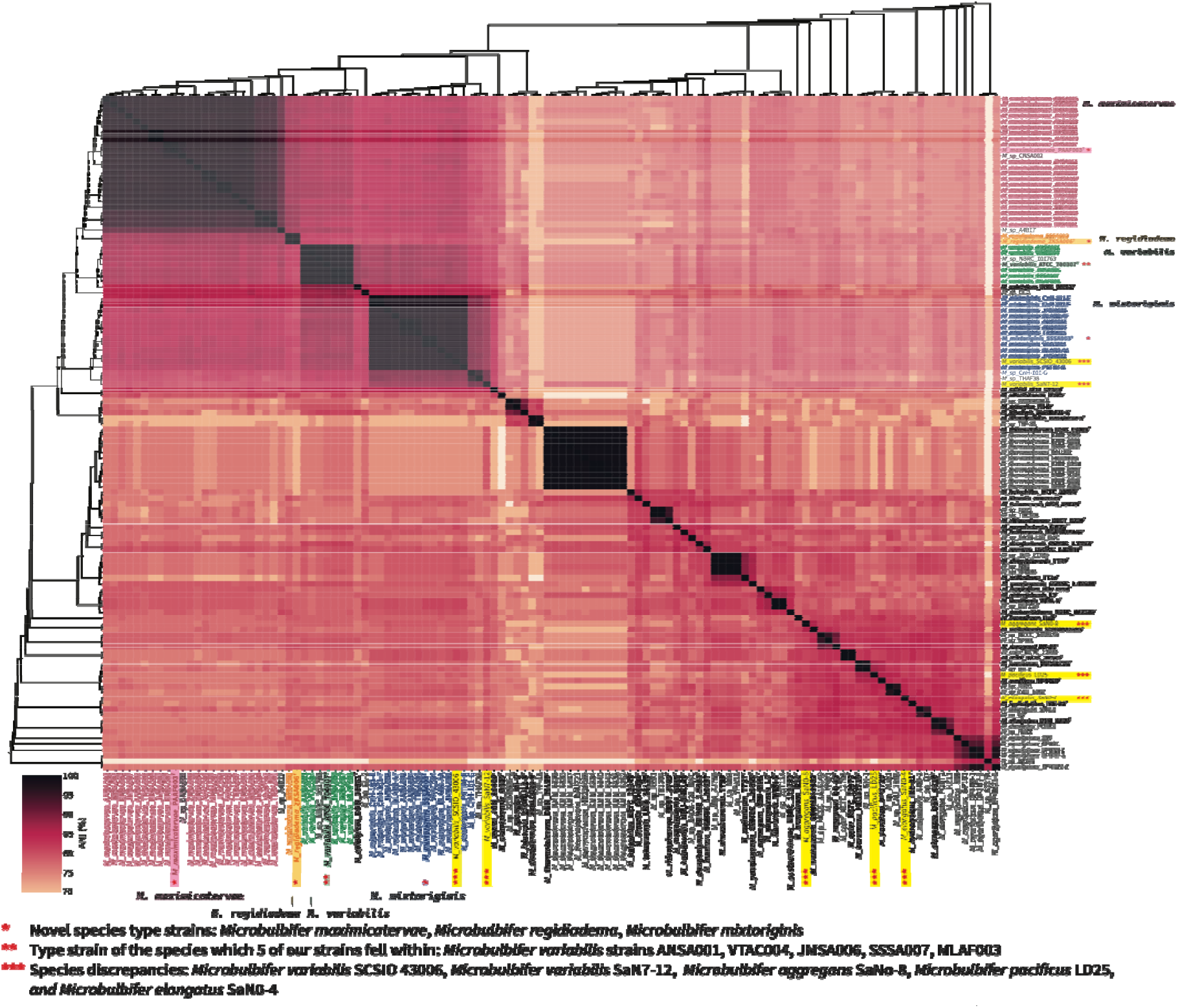
A genome similarity matrix illustrated as a heatmap colored according to ANI expressed as percentage values. The phylogenetic trees on both axes were reconstructed with the neighborhood joining method using pairwise distances calculated from ANI similarity scores. The novel species identified in this study are highlighted with color on the top right and bottom axes. Type strains of the newly proposed novel species are emphasized with one asterisk. Previously characterized type strain which some of our strains fell into is emphasized with two asterisks. Species discrepancies are emphasized with three asterisks.

GS-I: *Microbulbifer maximicatervae*

GS-II: *Microbulbifer regidiadema*

GS-III: *Microbulbifer mixtioriginis*

We propose *Microbulbifer maximicatervae* as the name for GS-I since the majority (61%) of *Microbulbifer* strains isolated from marine sponges were placed in this clade. Strain *M. maximicatervae* PAAF003^T^ was chosen as the type strain for this species. We propose *Microbulbifer regidiadema* as the name for GS-II bacteria due to the characteristic purple color of these microbes with strain *M. regidiadema* ZKSA006^T^ selected as the type strain. Finally, we propose *Microbulbifer mixtoriginis* as the name for GS-III because strains in this group were isolated from sponges as well as corals; *M. mixtoriginis* SSSA003^T^ was chosen as the type strain.

The ANI-based high-resolution phylogenomic organization for the *Microbulbifer* genus also allows us to propose several re-clarifications regarding previous species assignments. Several strains that were isolated from marine sponges and designated as GS-IV in our previous report grouped with the type strain *M. variabilis* ATCC 700307^T^ and have been now designated as such (Figure 1). The *Microbulbifer* strain CnH-101-G was previously grouped into GS-III. However, its ANI score to the GS-III type strain *M. mixtoriginis* SSSA003^T^ (*vide supra*) is only 89.7%. Therefore, strain CnH-101-G fails should not be classified as *Microbulbifer mixtoriginis* and we suggest for it to continue to be designated as *M*. sp. CnH-101-G, perhaps warranting a distinct species designation upon further examination in the future[15].

It was interesting to note that strains *M. variabilis* SCSIO 43006 and *M. variabilis* SaN7-12 did not group with the type strain *M. variabilis* ATCC 700307^T^ (ANI scores 87.3% and 82.9%, respectively) (Figure 1). Instead, *M. variabilis* SCSIO 43006 grouped closer to *M. mixtoriginis* SSSA003^T^ (ANI score 97.9%), and thus presents a case for potential taxonomic reassignment. In the same fashion, the strains *M. aggregans* SaN0-8, *M. pacificus* LD25, and *M. elongatus* SaN0-4 showed only modest ANI scores to their corresponding type strains (78.9% to *M. aggregans* CCB-MM1^T^, 85.6% to *M. pacificus* SPO729^T^, and 84.6% to *M. elongatus* DSM 6810^T^, respectively). This is not the first instance of high-resolution microbial whole-genome relatedness assessment revealing discrepancies in taxonomic classification; for example, a previous study for the *Bacillus* genus revealed similar disagreements [16]. Among the different classification strategies, ANI is gaining prominence in achieving the requisite taxonomic resolution for robust species designation [17]. For instance, a recent study found the ANI similarity threshold of 96.7% to be the preferred method for species designation in the *Streptomyces* genus [18]. Similarly, a 98% ANI similarity threshold was required to obtain an clear species demarcation for poxviruses [19].

The type strains for the three novel *Microbulbifer* species named above, along with the GS-IV *M. variabilis* strain VTAC004 were morphologically characterized based on epifluorescence microscopy (Figure 2A–2H). Strains *M. maximicatervae* PAAF003^T^, *M. mixtoriginis* SSSA003^T^, and *M. variabilis* VTAC004 present a cocci-shaped morphology with an opaque yellowish hue (Figure 2A, 2C–2E, 2G– 2H). On the other hand, *M. regidiadema* ZKSA006^T^ exhibited a purple, bacilli-shaped morphology (Figure 2B, 2F).

**Figure 2.**
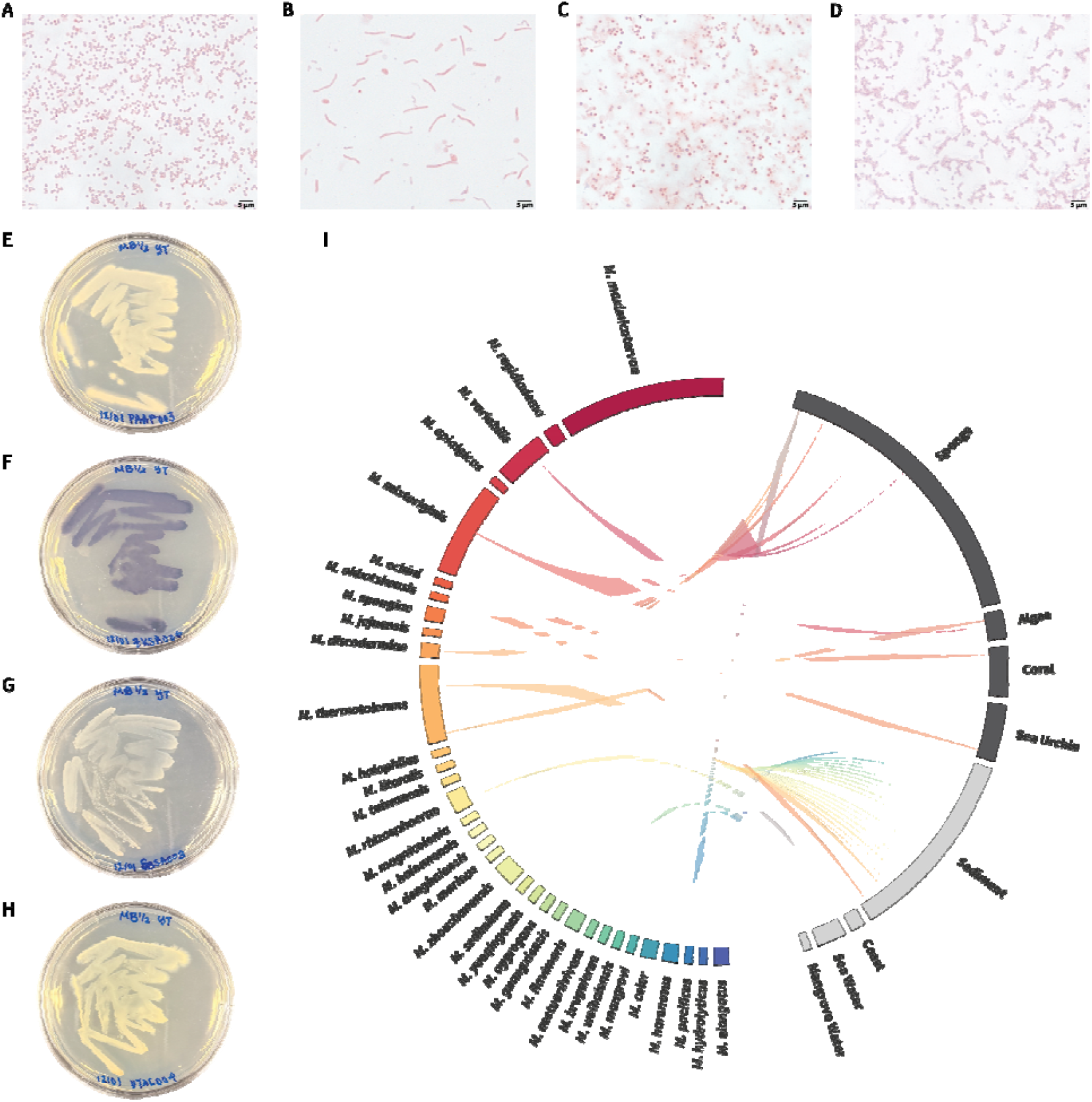
**(A–D)** High resolution Gram-stained images for *M. maximicatervae* PAAF003^T^, *M. regidiadema* ZKSA006^T^, *M. mixtoriginis* SSSA003^T^, and *M. variabilis* VTAC004, respectively. **(E– H)** Single strain inoculation of *M. maximicatervae* PAAF003^T^, *M. regidiadema* ZKSA006^T^, *M. mixtoriginis* SSSA003^T^, and *M. variabilis* VTAC004, respectively. **(I)** Circos diagram showing the relationship between each strain and their isolation source. The ANI-based species assignment for the strains is denoted. Species are arranged via the ANI phylogenetic order from Figure 1. Isolation sources were extrapolated via the available metadata for each genome. Eukaryotic sources are colored dark gray while microbiome-derived sources are colored light gray.

Next, we mapped the isolation sites of known *Microbulbifer* species on a world map (Figure S6, Dataset D2). This analysis established the cosmopolitan nature of the *Microbulbifer* genus in that it shares the same global dispersion patterns as other genera in the parent phylum *Pseudomonodota* such as *Pseudomonas, Pseudoalteromonas*, and *Vibrio* [20-22]. All 42 Microbulbifer strains isolated in our previous study originated from a single geographical site—Florida Keys, Florida, USA—likely leading to low species diversity. In contrast, *Microbulbifer* strains described by other studies have been distributed throughout the world and are associated with numerous holobionts likely resulting in higher species diversity.

In addition to sediments and the water column, *Microbulbifer* strains routinely make up the commensal microbiomes of marine invertebrates and macroalgae wherein they chemically interact and compete with other widely distributed bacterial genera [23]. To query the relationship between strain phylogeny and isolation sources, we inspected the metadata associated with each strain, extrapolating a source attribute through a combination of “isolation source”, “isolation site”, and “host” fields in the GenBank database. Additionally, we paired the metadata with the ANI-derived species at an ANI score threshold of 95% for species cutoff. With the species displayed in the order of their ANI-based phylogeny, a division between strains isolated from marine holobionts versus strains isolated from sediment and the water column was apparent (Figure 2I). Holobiont sources, such as sponges, algae, coral, and sea urchin have yielded 62 *Microbulbifer* strains organized into only 10 different species. On the other hand, sediment and the water column have yielded a much greater phylogenetic diversity, with 30 strains organized into 22 different species. Notable outliers were *M. okhotskensis* which was isolated from sediment but taxonomically grouped proximal to ten other holobiont-derived species and *M. pacificus*, which was conversely isolated from a sponge but was taxonomically grouped with the 23 other sediment/water-derived species (Figure 2I). As the *Microbulbifer* strain inventory expands, each strain adds a rich set of information not limited to its genomic content but expanding the available information for the ecological breadth of the genus based on where a strain comes from, how it was collected, and even the broad scale collection design. These attributes have facilitated the first ever meta-analysis of the *Microbulbifer* genus.

We already know that species-specific natural product biosynthetic potential contributes to chemical diversity within the *Microbulbifer* genus [10]. With a firm taxonomic profile for the genus now in hand, the stage is set for a more in-depth genomic analysis to discern the causal genetic differences that drive species divergence. *Microbulbifer* strains are ubiquitously detected as members of commensal microbiomes of marine holobionts wherein they interact not only with the host, but also with other prokaryotic community members. Interrogating whether these interactions are species specific, or not, will also be facilitated by the now benchmarked *Microbulbifer* taxonomy.

## Description of *Microbulbifer maximicatervae* sp. nov

*Microbulbifer maximicatervae* (ma.xi.mi.ca.ter’vae L. masc. adj. *maximus*, greatest, largest; L. fem. n. *caterva*, a crowd, a cluster; N.L. gen. n. *maximicatervae*, of the largest group). Cells grown on half strength artificial seawater media are an opaque yellowish hue with a cocci-shaped morphology, of about 0.7 µm length and 0.7 µm width. Members of the species are aerobes, growing at a pH optimum of 7.8, and a salinity optimum of 225 mM of NaCl at a temperature optimum of 30 ºC. Genome size is ∼5.2 Mbp with a G+C% content of 48.7.

The strain designated as type is PAAF003^T^ which was isolated from the commensal microbiome of the marine sponge *Aplysina fulva* collected in Florida Keys, Florida, USA (NCBI accession NZ_CP174319.1) [24].

### Description of *Micorbulbifer regidiadema* sp. nov

*Microbulbifer regidiadema* (re.gi.di.a.de’ma L. masc. adj. *regius*, royal; L. fem. n. *diadema*, crown, diadem; N.L. fem. n. *regidiadema*, royal crown). Cells grown on half strength artificial seawater media are purple with a bacilli-shaped morphology, of about 7.0 µm in length and 0.7 µm width. Members of the species are aerobes, growing at a pH optimum of 7.8, and a salinity optimum of 225 mM NaCl at a temperature optimum of 30 ºC. Genome size is ∼4.9 Mbp with a G+C% content of 51.3.

The strain designated as type is ZKSA006^T^ was isolated from the commensal microbiome of the marine sponge *Smenospongia aurea* collected in Florida Keys, Florida, USA (NCBI accession NZ_CP174297.1) [24].

### Description of *Micorbulbifer mixtoriginis* sp. nov

*Microbulbifer mixtoriginis* (mix.to.ri.gi’nis L. fem. pres. part. mixta, mixed; L. fem. n. origo, origin; N.L. gen. n. mixtoriginis, of mixed origin). Cells grown on half strkey weength artificial seawater media are an opaque yellowish hue with a cocci-shaped morphology, of about 0.7 µm length and 0.6 µm width. Members of the species are aerobes, growing at a pH optimum of 7.8, and a salinity optimum of 225 mM NaCl at a temperature optimum of 30 ºC. Genome size is ∼4.8 Mbp with a G+C% content of 49.3.

The strain designated as type is SSSA003^T^ was isolated from the commensal microbiomes of the marine sponge *S. aurea* collected in Florida Keys, Florida, USA (NCBI accession NZ_CP174314.1) [24].

## Supporting Information

Supporting Information document accompanying this article containing detailed description of Materials and Methods, Supplementary Table S1, Supplementary Figures S1–S6, and Supplementary References. Additional Supplementary Datasets D1–D2 are provided as spreadsheets.

Supplementary Dataset D1: ANI scores for *Microbulbifer* strains.

Supplementary Dataset D2: Isolation source of *Microbulbifer* strains.

## Data Deposition

The three new type strains PAAF003^T^, ZKSA006^T^, and SSS003^T^ are registered under the SeqCode Registry (seqco.de/r:3zzf79a7) [14].

## Acknowledgements

Permits for coral collection were received from the Florida Keys National Marine Sanctuary (Permit Nos. FKNMS-2017-128-A2 and FKNMS-2019-160).

