## supplement_document for "Phylogenomic description of three novel species of the *Microbulbifer* genus, phylum Pseudomonadota, isolated from marine sponges and corals"

**Supplementary Materials and Methods**

**Phylogenetic tree construction**

16S rRNA sequence coordinates were extracted with Barrnap 0.9, sequences extracted with BEDTools and aligned with Mafft version 7 [1-3]. The phylogenetic tree was constructed with IQ-TREE3 [4]. The analysis was run on 4 threads (-*nt 4*) with automatic ModelFinder (-*m MFP*). Branch support was assessed through 1,000 ultrafast bootstrap replicates (-*B 1000*) along with 1,000 replicates of the Shimodaira-Hasegawa approximate Likelihood Ratio Test (SH-aLRT) (-*alrt 1000*) [5-8]. Outgroups were specified as *Aeromonas hydrophila* ATCC 7966, *Escherichia coli* ATCC 11775, and *Vibrio cholerae* ATCC 14035 (*-o A hydrophila ATCC 7966, E coli ATCC 11775, V cholerae ATCC 14035*) [9]. The consensus phylogeny was visualized with FigTree v1.4.4 [10]. An automated MLSA pipeline, automlsa2, was employed to extract, align, and concatenate 15 housekeeping genes and generate a maximum likelihood tree [11]. The consensus phylogeny was visualized with FigTree v1.4.4 [10].

**Average Nucleotide Identity (ANI) heatmap representation**

Pangenome was created using PanTools [12]. ANI similarity indices of the pangenome were calculated using FastANI and MASH to extract orthologous genes and produce a Rscript which constructed a neighborhood joining tree [13,14]. Heatmap representation of the pairwise similarity index was visualized using seaborn [15]. The consensus phylogeny was visualized with FigTree v1.4.4 [10].

**Gram staining and strain imaging**

From glycerol stocks of *Microbulbifer* strains stored at –80 °C were inoculated onto agar plates containing half strength marine broth (MB ½) and incubated at 30 °C for 5 d. An inoculation loop was used to transfer a single drop of water onto a microscope slide. A minimal amount of the bacterial colony was transferred from the agar plate onto the droplet of water on the microscopic slide. Once the droplet has air dried, use heat to fix the cells to the glass slide. Add two to three droplets of crystal violet stain over the fixed culture allowing stain to rest for 60 seconds. Rinse slides with water to wash off excess crystal violet stain. Add two to three droplets of iodine solution over the fixed culture allowing solution to rest for 60 seconds. Rinse slides with water to wash off excess iodine solution. Hold microscope slide at a 45° angle and add two to three drops of decolorizer. Add two to three drops of safranin counterstain over the fixed culture allowing stain to rest for 60 seconds. Rinse slides with water to wash off excess safranin. Microscope slides were imaged using the Zeiss Axio Observer Z1 color/fluorescent microscope in brightfield lighting using a 100× oil immersion objective lens.

**Geography bubble map plot**

The geography bubble map was created by assigning a latitude and longitude to each strain collected inferred from the collection location metadata attribute. This data was collated into a comma-separated-values file which was then transformed and cleaned before serving as the input for the plotly geographic visualization [16]. Certain datapoints were “jittered” for visual clarity.

**Circos isolation source plot**

The Circos plot was created by manually ascertaining the isolation source from the available metadata attributes [17]. This data was put into a comma-separated-values file transformed and cleaned, removing all strains that weren’t assigned a species, before inputted into the pycircilize draw function. Of note the order of species on the left-hand side are static to reflect the phylogenetic order whereas the order of isolation sources is inferred by pycircilize to maximize visual clarity.

**Supplementary Tables**

**Table S1:** *Microbulbifer* strains used in this study

| **Strain name** | **Source** | **Genome Size** | **GC Content (%)** | **References** |
| --- | --- | --- | --- | --- |
| **Sponge-derived *Microbulbifer* strains with genomes sequenced from our study** | | | | |
| *Microbulbifer* sp. ANSA001 | *Smenospongia aurea* | 5.0 Mb | 48.83 | Genbank [18] |
| *Microbulbifer* sp. ANSA002 | *Smenospongia aurea* | 4.8 Mb | 49.27 | Genbank [18] |
| *Microbulbifer* sp. ANSA003 | *Smenospongia aurea* | 5.6 Mb | 48.83 | Genbank [18] |
| *Microbulbifer* sp. ANSA004 | *Smenospongia aurea* | 4.8 Mb | 49.27 | Genbank [18] |
| *Microbulbifer* sp. ANSA005 | *Smenospongia aurea* | 5.2 Mb | 48.69 | Genbank [18] |
| *Microbulbifer* sp. AVAC002 | *Aiolochroia crassa* | 5.2 Mb | 48.71 | Genbank [18] |
| *Microbulbifer* sp. EKSA005 | *Smenospongia aurea* | 5.3 Mb | 48.56 | Genbank [18] |
| *Microbulbifer* sp. EKSA006 | *Smenospongia aurea* | 5.2 Mb | 48.62 | Genbank [18] |
| *Microbulbifer* sp. EKSA007 | *Smenospongia aurea* | 5.3 Mb | 48.63 | Genbank [18] |
| *Microbulbifer* sp. EKSA008 | *Smenospongia aurea* | 5.2 Mb | 48.71 | Genbank [18] |
| *Microbulbifer* sp. JMSA002 | *Smenospongia aurea* | 4.8 Mb | 49.33 | Genbank [18] |
| *Microbulbifer* sp. JMSA003 | *Smenospongia aurea* | 5.2 Mb | 48.62 | Genbank [18] |
| *Microbulbifer* sp. JMSA004 | *Smenospongia aurea* | 5.2 Mb | 48.62 | Genbank [18] |
| *Microbulbifer* sp. JMSA006 | *Smenospongia aurea* | 5.0 Mb | 48.73 | Genbank [18] |
| *Microbulbifer* sp. JMSA007 | *Smenospongia aurea* | 5.2 Mb | 48.53 | Genbank [18] |
| *Microbulbifer* sp. JMSA008 | *Smenospongia aurea* | 4.8 Mb | 49.34 | Genbank [18] |
| *Microbulbifer* sp. JTAC008 | *Aiolochroia crassa* | 5.3 Mb | 48.64 | Genbank [18] |
| *Microbulbifer* sp. PAAF003 | *Aplysina fulva* | 5.2 Mb | 48.71 | Genbank [18] |
| *Microbulbifer* sp. SSSA002 | *Smenospongia aurea* | 5.1 Mb | 51.45 | Genbank [18] |
| *Microbulbifer* sp. SSSA003 | *Smenospongia aurea* | 4.8 Mb | 49.34 | Genbank [18] |
| *Microbulbifer* sp. SSSA005 | *Smenospongia aurea* | 5.4 Mb | 48.63 | Genbank [18] |
| *Microbulbifer* sp. SSSA007 | *Smenospongia aurea* | 5.0 Mb | 48.73 | Genbank [18] |
| *Microbulbifer* sp. SSSA008 | *Smenospongia aurea* | 5.2 Mb | 48.81 | Genbank [18] |
| *Microbulbifer* sp. TRSA001 | *Smenospongia aurea* | 5.3 Mb | 48.70 | Genbank [18] |
| *Microbulbifer* sp. TRSA002 | *Smenospongia aurea* | 5.5 Mb | 48.61 | Genbank [18] |
| *Microbulbifer* sp. TRSA005 | *Smenospongia aurea* | 4.8 Mb | 49.25 | Genbank [18] |
| *Microbulbifer* sp. TRSA007 | *Smenospongia aurea* | 5.1 Mb | 48.82 | Genbank [18] |
| *Microbulbifer* sp. VAAC004 | *Aiolochroia crassa* | 4.8 Mb | 49.34 | Genbank [18] |
| *Microbulbifer* sp. VTAC004 | *Aiolochroia crassa* | 4.9 Mb | 48.87 | Genbank [18] |
| *Microbulbifer* sp. VVAC002 | *Aiolochroia crassa* | 5.3 Mb | 48.71 | Genbank [18] |
| *Microbulbifer* sp. ZKSA002 | *Smenospongia aurea* | 5.2 Mb | 48.72 | Genbank [18] |
| *Microbulbifer* sp. ZKSA004 | *Smenospongia aurea* | 5.2 Mb | 48.62 | Genbank [18] |
| *Microbulbifer* sp. ZKSA006 | *Smenospongia aurea* | 4.9 Mb | 51.32 | Genbank [18] |
| *Microbulbifer* sp. MLAF003 | *Aplysina fulva* | 4.9 Mb | 49.00 | Genbank [18] |
| *Microbulbifer* sp. VAAF005 | *Aplysina fulva* | 5.4 Mb | 48.50 | Genbank [18] |
| *Microbulbifer* sp. MKSA007 | *Smenospongia aurea* | 5.8 Mb | 49.00 | Genbank [18] |
| **Coral-derived *Microbulbifer* strains with genomes sequenced from our study** | | | | |
| *Microbulbifer* sp. CnH-101-E | *Colpophyllia natans* | 5.0 Mb | 49.24 | Genbank [18] |
| *Microbulbifer* sp. CnH-101-F | *Colpophyllia natans* | 5.0 Mb | 49.24 | Genbank [18] |
| *Microbulbifer* sp. CnH-101-G | *Colpophyllia natans* | 4.6 Mb | 50.21 | Genbank [18] |
| *Microbulbifer* sp. DLAB2-AA | *Diploria labyrinthiformis* | 4.9 Mb | 49.34 | Genbank [18] |
| *Microbulbifer* sp. DLAB2-AF | *Diploria labyrinthiformis* | 4.8 Mb | 49.36 | Genbank [18] |
| *Microbulbifer* sp. PSTR4-B | *Pseudodiploria (*formerly *Diploria) strigosa* | 5.0 Mb | 49.42 | Genbank [18] |
| ***Microbulbifer* type strains with available draft genomes** | | | | |
| *Microbulbifer aestuariivivens* NBRC 112533^T^ | Tidal flat sediment | 3.4 Mb | 59.50 | Genbank [19] |
| *Microbulbifer* *aggregans* CBB-MM1^T^ | Mangrove sediment | 3.9 Mb | 59.00 | Genbank [20] |
| *Microbulbifer bruguierae* H12^T^ | Mangrove plant | 4.5 Mb | 56.50 | Genbank [21] |
| *Microbulbifer celer* KCTC 12973^T^ | Marine solar saltern | 4.3 Mb | 57.00 | Genbank [22] |
| *Microbulbifer donghaiensis* CGMCC 1.7063^T^ | Marine sediment | 4.3 Mb | 59.50 | Genbank [23] |
| *Microbulbifer discodermiae 2201CG32-9*^T^ | Marine sponge | 4.6 Mb | 56.50 | Genbank [24] |
| *Microbulbifer echini* JCM 30400^T^ | Sea urchin | 4.4 Mb | 51.00 | Genbank [25] |
| *Microbulbifer elongatus* DSM 6810^T^ | Sea water | 4.2 Mb | 57.50 | Genbank ^[26]^ |
| *Microbulbifer epialgicus* DSM 18651^T^ | Marine algae | 5.7 Mb | 49.00 | Genbank [27] |
| *Microbulbifer flavimaris* WRN-8^T^ | Marine sediment | 3.6 Mb | 60.00 | Genbank [28] |
| *Microbulbifer guangxiensis* L3^T^ | Tidal flat sediment | 3.7 Mb | 60.00 | Genbank [29] |
| *Microbulbifer halophilus* KCTC 12848^T^ | Saline soil | 4.7 Mb | 61.50 | Genbank [30] |
| *Microbulbifer harenosus* HB161719^T^ | Coastal sand | 4.7 Mb | 58.00 | Genbank [31] |
| *Microbulbifer hydrolyticus* IRE-31^T^ | Black liquor | 4.2 Mb | 57.50 | Genbank [32] |
| *Microbulbifer jejuensis* 2304DJ12-6^T^ | Marine sponge | 4.7 Mb | 53.50 | Genbank [24] |
| *Microbulbifer magnicolonia* GG15^T^ | Tidal flat sediment | 4.3 Mb | 61.50 | Genbank [33] |
| *Microbulbifer mangrovi* DD-13^T^ | Mangrove water | 4.5 Mb | 57.00 | Genbank [34] |
| *Microbulbifer marinus* CGMCC 1.10657^T^ | Marine sediment | 4.0 Mb | 60.00 | Genbank [35] |
| *Microbulbifer pacificus* SPO729^T^ | Marine sponge | 4.2 Mb | 58.50 | Genbank [36] |
| *Microbulbifer rhizosphaerae* CECT 8799 ^T^ | Plant | 5.3 Mb | 60.00 | Genbank [37] |
| *Microbulbifer sediminum* TT37^T^ | Tidal flat sediment | 3.9 Mb | 61.00 | Genbank [29] |
| *Microbulbifer spongiae* MI-G^T^ | Marine sponge | 4.5 Mb | 53.50 | Genbank [38] |
| *Microbulbifer taiwanensis* LMG 26125^T^ | Coastal soil | 4.8 Mb | 60.00 | Genbank [39] |
| *Microbulbifer thermotolerans* DSM 19189^T^ | Deep-sea sediment | 3.9 Mb | 56.90 | Genbank [40] |
| *Microbulbifer variabilis* ATCC 700307^T^ | Marine algae | 4.8 Mb | 49.00 | Genbank [27] |
| *Microbulbifer yueqingensis* CGMCC 1.10658^T^ | Marine sediment | 3.7 Mb | 62.00 | Genbank [35] |
| *Microbulbifer zhoushanensis* TT30^T^ | Tidal flat sediment | 4.1 Mb | 61.50 | Genbank [29] |
| ***Microbulbifer* type strains with only 16S rRNA gene sequences available** | | | | |
| *Microbulbifer arenaceous* RSBr-1T^T^ | Red Sandstone | n/a | n/a | Genbank [41] |
| *Microbulbifer agarlyticus* JAMB A3^T^ | Deep-sea sediment | n/a | n/a | Genbank [40] |
| *Microbulbifer chitinlyticus* JCM 15148^T^ | Mangrove forests | n/a | n/a | Genbank [42] |
| *Microbulbifer gwangyangensis* GY2^T^ | Tidal flat | n/a | n/a | Genbank [36] |
| *Microbulbifer maritimus* TF-17^T^ | Intertidal sediment | n/a | n/a | Genbank [43] |
| *Microbulbifer okhotskensis* KMM 9862^T^ | Okhotsk Sea sediment | n/a | n/a | Genbank [44] |
| *Microbulbifer okinawensis* ABABA23^T^ | Mangrove forests | n/a | n/a | Genbank [42] |
| **Outgroup strains with available draft genomes** | | | | |
| *Aeromonas hydrophila* ATCC 7966^T^ | Water | 4.7 Mb | 61.50 | Genbank [45] |
| *Escherichia coli* ATCC 11775^T^ | Digestive tract | 5.0 Mb | 50.50 | Genbank [46] |
| *Vibrio cholerae* ATCC 14035^T^ | Human | 4.0 Mb | 47.50 | Genbank [47] |
| ***Microbulbifer* strains with available draft genomes** | | | | |
| *Microbulbifer* sp. 2205BS26-8 | Marine sponge | 3.9 Mb | 54.00 | n/a |
| *Microbulbifer* sp. A4B17 | Sea water | 5.0 Mb | 48.50 | Genbank [48] |
| *Microbulbifer* sp. ALW1 | Kelp | 4.7 Mb | 57.00 | Genbank [49] |
| *Microbulbifer* sp. CAU 1566 | Soil | 4.4 Mb | 57.00 | n/a |
| *Microbulbifer* sp. CNSA002 | Marine sponge | 5.3 Mb | 48.50 | n/a |
| *Microbulbifer* sp. GL-2 | Marine fish intestine | 5.0 Mb | 49.50 | Genbank [50] |
| *Microbulbifer* sp. HZ11 | Sea water | 4.2 Mb | 56.50 | Genbank [51] |
| *Microbulbifer* sp. JSM ZJ756 | Tidal zone sediment | 4.2 Mb | 61.00 | n/a |
| *Microbulbifer* sp. M83 | Tidal flat sediment | 4.2 Mb | 61.00 | n/a |
| *Microbulbifer* sp. MCCC 1A16149 | Sediment | 4.5 Mb | 57.50 | n/a |
| *Microbulbifer* sp. NBRC 101763 | Algae | 4.9 Mb | 49.00 | n/a |
| *Microbulbifer* sp. Q7 | Sea cucumber | 4.0 Mb | 58.50 | Genbank [52] |
| *Microbulbifer* sp. RZ01 | Coastal environment | 5.4 Mb | 60.00 | Genbank [53] |
| *Microbulbifer* sp. S227A | Brittle star | 5.7 Mb | 62.00 | n/a |
| *Microbulbifer* sp. *SAOS-129* SWC | Mangrove sediment | 4.6 Mb | 61.00 | n/a |
| *Microbulbifer* sp. SH-1 | Coastal Soil | 4.7 Mb | 58.00 | n/a |
| *Microbulbifer* sp. TB1203 | Coastal environment | 5.4 Mb | 60.00 | Genbank [53] |
| *Microbulbifer* sp. THAF38 | Stony coral | 4.8 Mb | 50.00 | n/a |
| *Microbulbifer* sp. TYP-18 | Coral | 4.5 Mb | 56.50 | n/a |
| *Microbulbifer* sp. YPW1 | Mangrove sediment | 4.6 Mb | 57.50 | Genbank [54] |
| *Microbulbifer* sp. YPW16 | n/a | 4.2 Mb | 61.00 | n/a |
| *Microbulbifer* sp. ZGT114 | Erba brine-seawater | 3.6 Mb | 60.00 | n/a |
| *Microbulbifer* agarilyticus DP3N7-3 | Sediment | 4.2 Mb | 56.00 | n/a |
| *Microbulbifer* agarilyticus DP4N21-6 | Sediment | 4.1 Mb | 56.00 | n/a |
| *Microbulbifer agarilyticus* DP5N0-1 | Sediment | 4.1 Mb | 56.00 | n/a |
| *Microbulbifer agarilyticus* GP101 | Invertebrate gut wall | 4.3 Mb | 55.50 | n/a |
| *Microbulbifer agarilyticus* S89 | n/a | 3.9 Mb | 57.00 | n/a |
| *Microbulbifer aggregans* SaN0-8 | Mangrove sediment | 4.8 Mb | 56.00 | n/a |
| *Microbulbifer celer* KCTC 12793 | Marine solar saltern | 4.3 Mb | 57.50 | n/a |
| *Microbulbifer elongatus* PORT2 | Surface seawater | 4.2 Mb | 57.50 | Genbank [55] |
| *Microbulbifer elongatus* SaN0-4 | Mangrove sediment | 4.2 Mb | 57.00 | n/a |
| *Microbulbifer pacificus* LD25 | Salt water hot spring | 4.3 Mb | 57.50 | Genbank [56] |
| *Microbulbifer salipaludis* SN0-2 | Offshore sediment | 4.0 Mb | 58.50 | n/a |
| *Microbulbifer thermotolerans* DAU221 | Marine sediment | 3.9 Mb | 56.50 | Genbank [57] |
| *Microbulbifer thermotolerans* HB226069 | Brown macroalgae | 4.0 Mb | 56.50 | Genbank [58] |
| *Microbulbifer thermotolerans* KMM 6242 | Purple sea urchin | 3.9 Mb | 56.50 | Genbank [59] |
| *Microbulbifer thermotolerans* KMM 6262 | Purple sea urchin | 3.9 Mb | 56.50 | Genbank [59] |
| *Microbulbifer thermotolerans* KMM 6834 | Purple sea urchin | 3.9 Mb | 56.50 | Genbank [59] |
| *Microbulbifer thermotolerans* KMM 6835 | Purple sea urchin | 3.9 Mb | 56.50 | Genbank [59] |
| *Microbulbifer thermotolerans* KMM 6836 | Purple sea urchin | 3.9 Mb | 56.50 | Genbank [59] |
| *Microbulbifer thermotolerans* KMM 6837 | Purple sea urchin | 4.0 Mb | 56.50 | Genbank [59] |
| *Microbulbifer thermotolerans* KMM 6838 | Purple sea urchin | 3.9 Mb | 56.50 | Genbank [59] |
| *Microbulbifer thermotolerans* KMM 6840 | Bottom sediment | 3.9 Mb | 56.50 | Genbank [59] |
| *Microbulbifer variabilis* SaN7-12 | Mangrove sediment | 4.8 Mb | 50.00 | n/a |
| *Microbulbifer variabilis* SCSIO 43006 | Stony coral | 4.9 Mb | 49.50 | n/a |

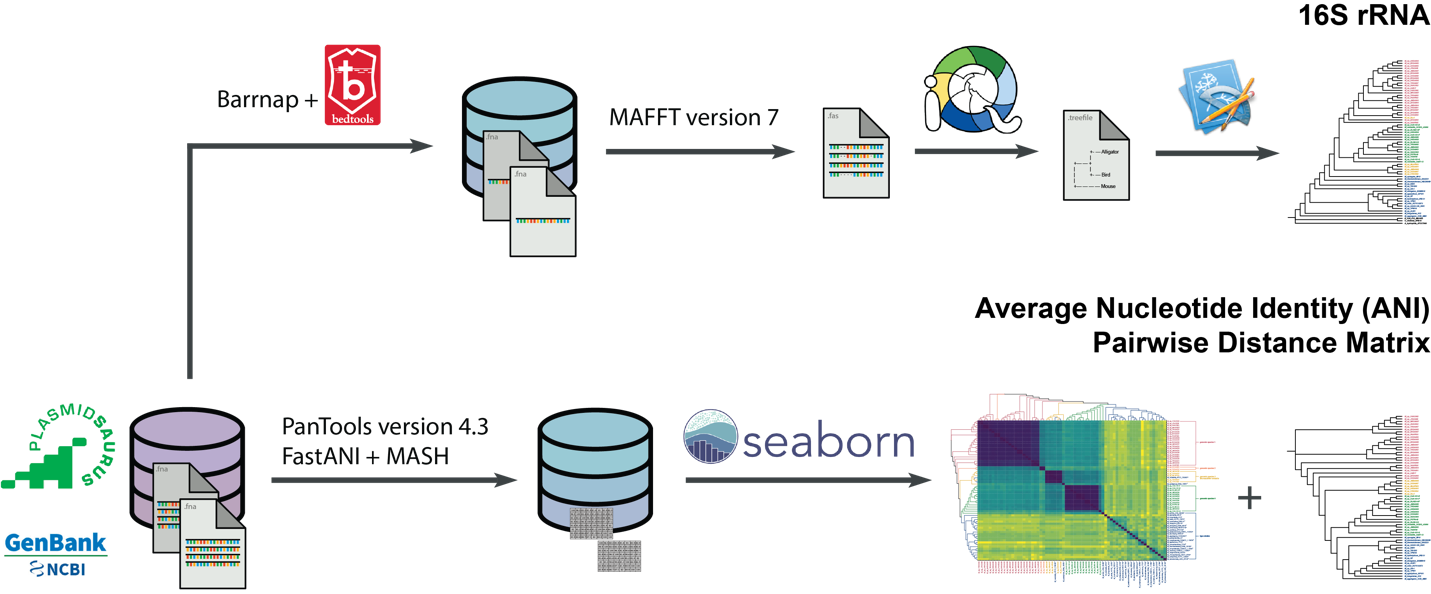

**Figure S1.** Cartoon depicting the workflow to assign a genus and putative species using 16S rRNA, orthologous genes, and housekeeping genes. Barrnap 0.9, Mafft version 7, and IQ-TREE3 were employed to extract 16S rRNA sequences and generate a maximum likelihood tree. PanTools, FastANI, and MASH were employed to extract orthologous genes and generate a neighborhood joining tree and heatmap representation of pairwise distances [1,3,4,12-14].

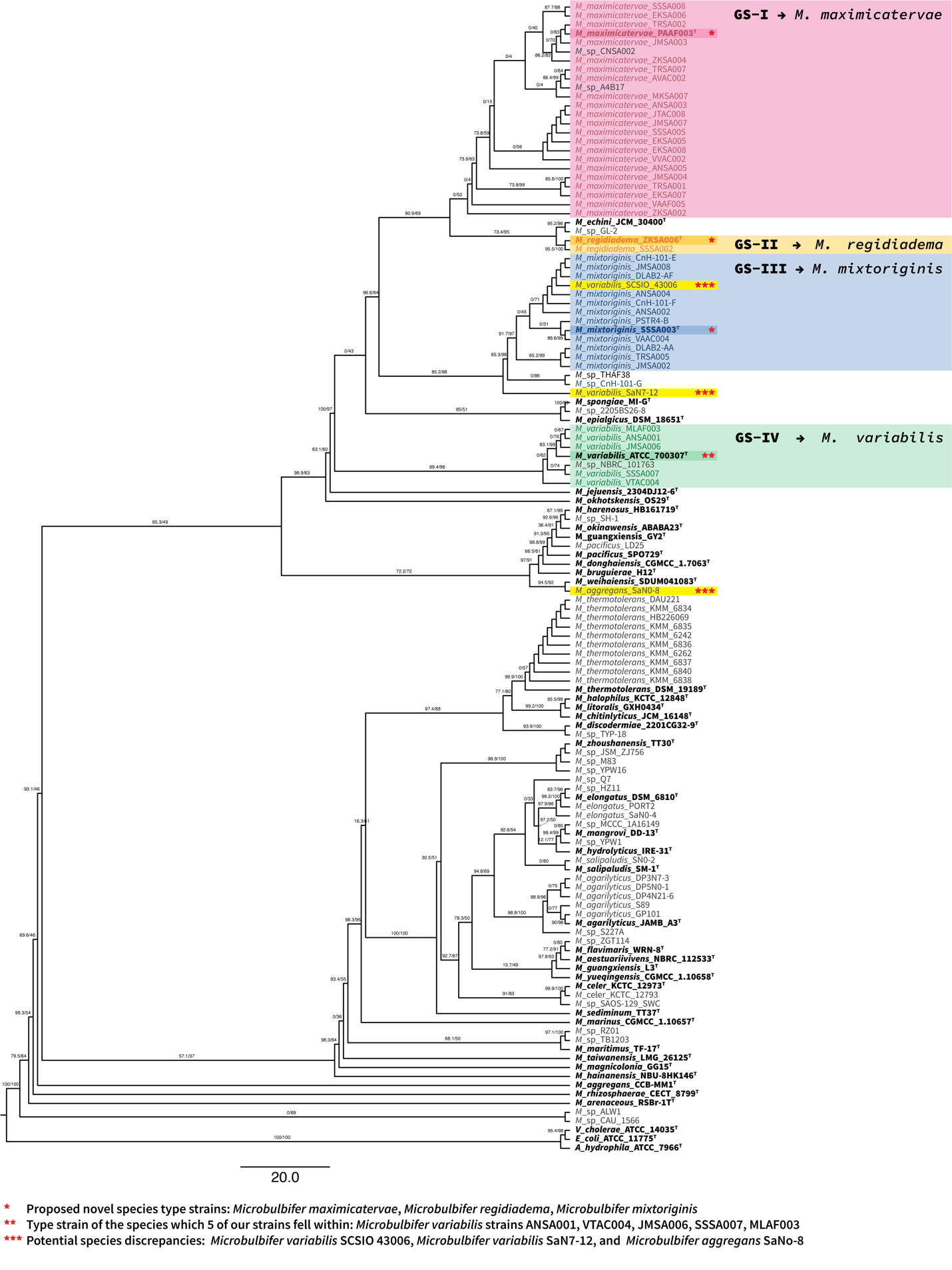

**Figure S2.** Phylogenetic tree reconstructed by the maximum-likelihood method using 16S rRNA. Subclades are highlighted to effectively compare phylogenetic estimations across three methods of phylogenetic tree construction. Three strains—*Escherichia coli K12 MG1655, Vibrio cholerae RFB16,* and *Aeromonas hydrophilia ATCC7966—*are selected as outgroups to root the tree. Shimodaira-Hasegawa approximate Likelihood Ratio Test (SH-aLRT)/ultrafast bootstrap values are both displayed [5,7,8].

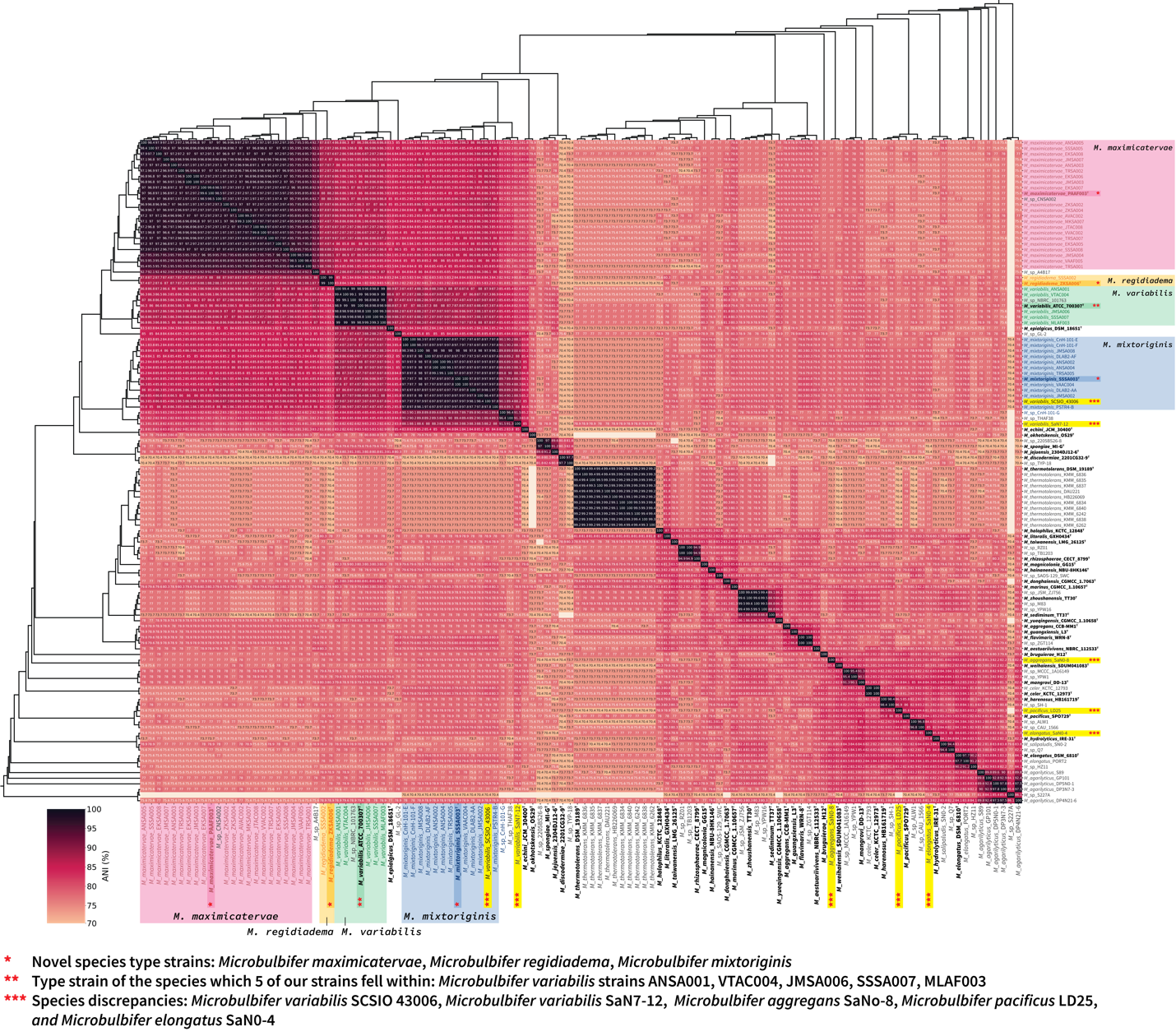

**Figure S3.** Heatmap representation of pairwise distances in a hierarchal clustering analysis derived from ANI based calculations. Clades with ANI similarity scores ≥95% formed genomic species (genomic species I– IV). Note an exception with this grouping can be found within genomic species III. *Microbulbifer* sp. CnH-101-G has ANIs score which are ≥89% relative to other strains found in genomic species III but are across the board ≤95%. Although an ANI similarly score of ≤95% denotes that strain CnH-101-G is likely not the same species as the rest of genomic species III, genomic species III is its closest related species.

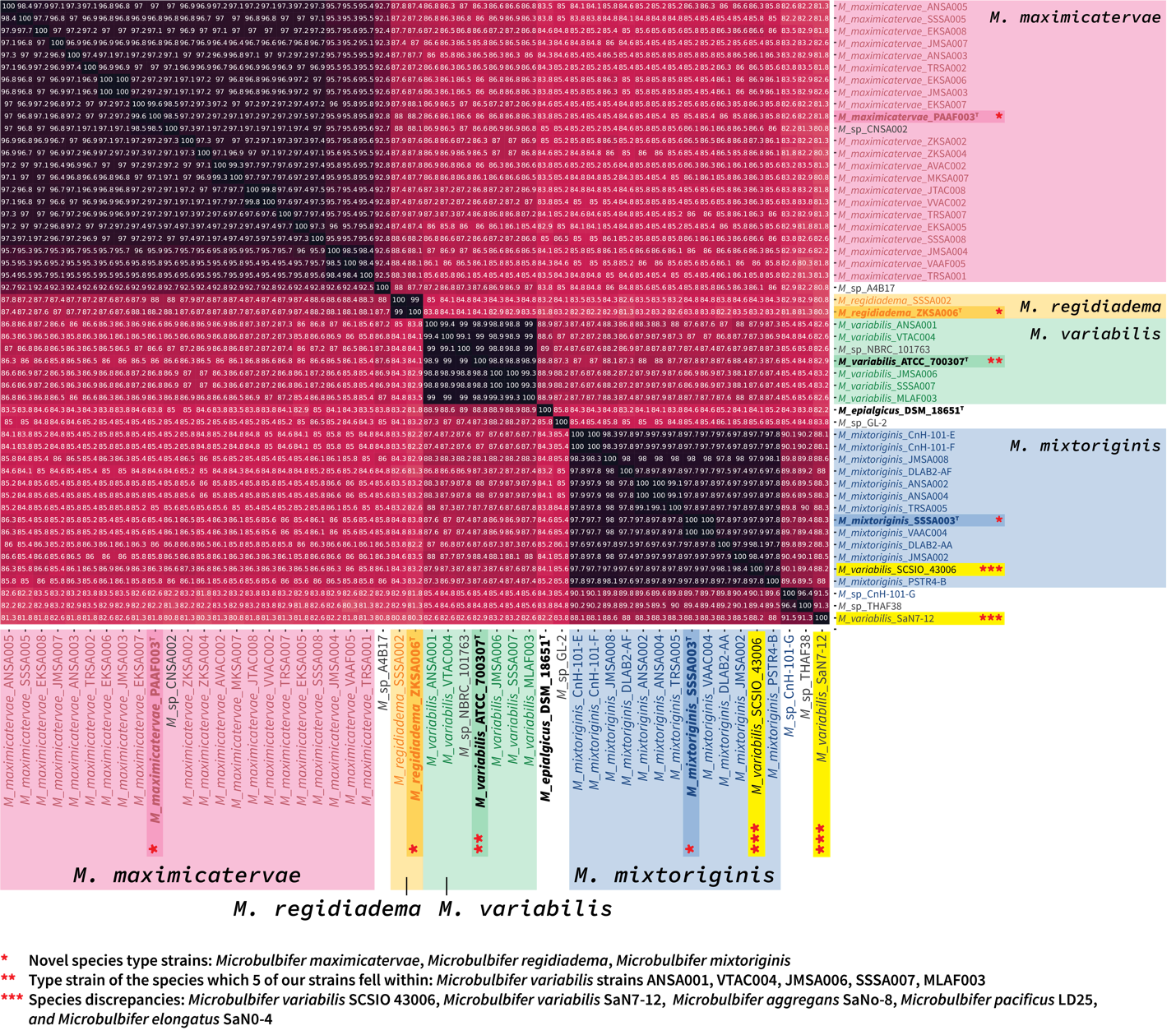

**Figure S4.** Expanded ANI to highlight the pairwise ANI scores across four species. A cutoff of 95% in ANI similarity score delineated a new species [60].

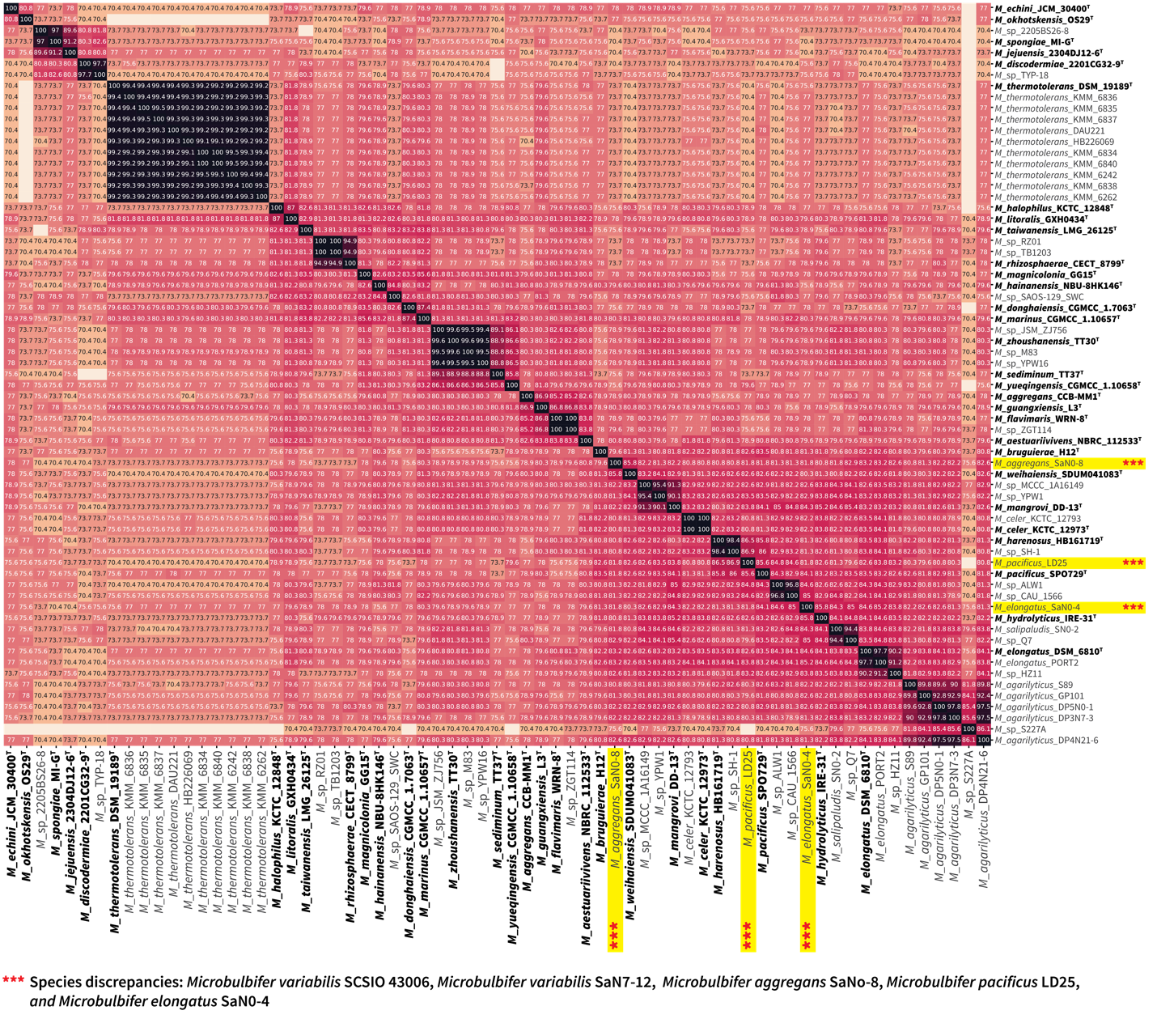

**Figure S5.** Expanded ANI to highlight the pairwise ANI scores across 29 species. A cutoff of 95% in ANI similarity score delineated a new species [60].

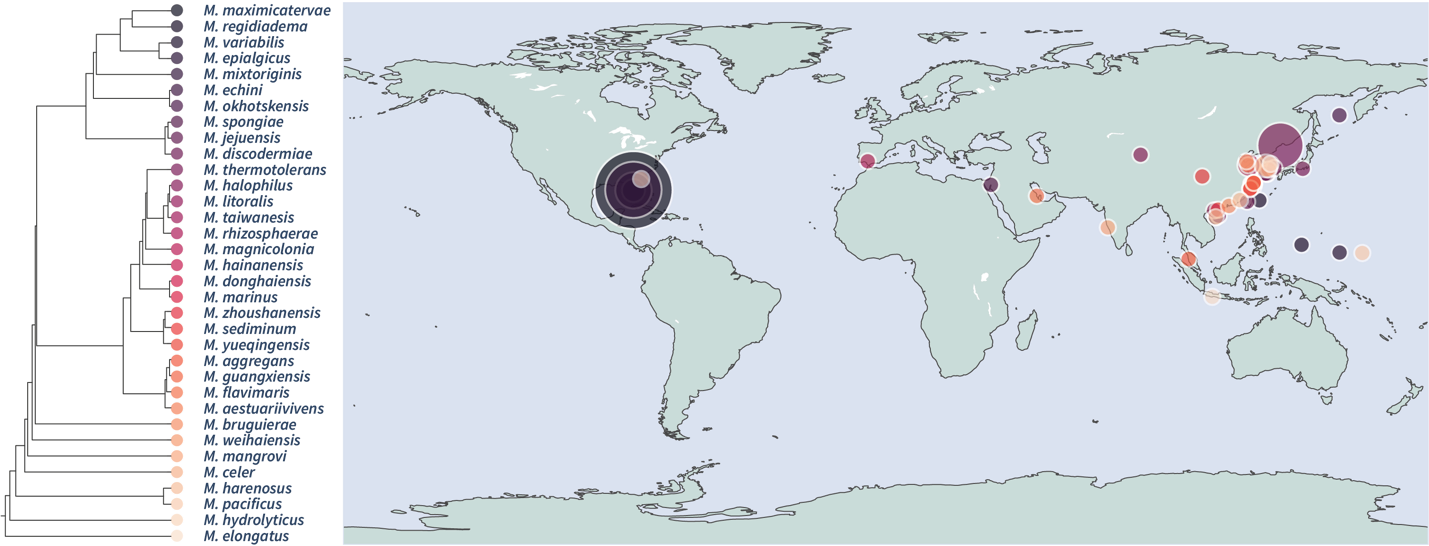

**Figure S6.** Geographical bubble map of all available species with location attached to their metadata. The left side phylogenetic tree is a reproduction of the ANI tree illustrated in Figure S3. The bubbles are represented by the color each species falls on the phylogenetic hierarchy denoted in the phylogenetic tree.
